# Linking genomic prediction for muscle fat content in Atlantic salmon to underlying changes in lipid metabolism regulation

**DOI:** 10.1101/2023.08.29.555338

**Authors:** Thomas N. Harvey, Hanne Dvergedal, Lars Grønvold, Yang Jin, Jørgen Ødegård, Sven Arild Korsvoll, Tim Knutsen, Torgeir R. Hvidsten, Simen R. Sandve

**Affiliations:** Department of Animal and Aquacultural Sciences, Faculty of Biosciences, Norwegian University of Life Sciences, P. O. Box 5003, NO-1433 Ås, Norway; AquaGen AS, P. O. Box 1240, NO-7462 Trondheim, Norway; Faculty of Chemistry, Biotechnology and Food Sciences, Norwegian University of Life Sciences, NO-1432 Ås, Norway

**Keywords:** Atlantic salmon, genomic selection, lipid metabolism, fat content, gene expression, eQTL

## Abstract

Muscle fat content is an important production trait in Atlantic salmon (*Salmo salar*) because it influences the flavor, texture, and nutritional properties of the fillet. Genomic selection can be applied to alter muscle fat content, however how such selection changes the underlying molecular physiology of these animals is unknown. Here, we examine the link between genomic prediction and underlying molecular physiology by correlating genomic breeding values for fat content to liver gene expression in 184 fish. We found that Salmon with higher genomic breeding values had higher expression of genes in lipid metabolism pathways. This included key lipid metabolism genes *hmgcrab*, *fasn-b*, *fads2d5*, and *fads2d6*, and lipid transporters *fatp2f*, *fabp7b*, and *apobc*. We also found several regulators of lipid metabolism with negative correlation to genomic breeding vales, including *pparg-b*, *fxr-a*, and *fxr-b*. A quantitative trait loci analysis for variation in gene expression levels (eQTLs) for 167 trait associated genes found that 71 genes had at least one eQTL, and that most were trans eQTLs. Closer examination revealed distinct eQTL clustering on chromosomes 3 and 6, indicating the presence of putative common regulator in these regions. Taken together, these results suggest that increased fat content in high genomic breeding value salmon is associated with elevated lipid synthesis, elevated lipid transport, and reduced glycerolipid breakdown; and that this is at least partly achieved by selection on genetic variants that impact the function of top-level transcription factors involved in liver metabolism. Our study sheds light on how genomic selection alters lipid content in Atlantic salmon, and the results could be used to prioritize SNPs to improve the efficiency of genomic selection in the future.

## 1 Background

Intramuscular fat content is an important quality parameter in most production animals because it influences the texture, flavor, and nutritional properties of the meat. The latter is especially important in Atlantic salmon as it is considered an excellent source of healthy omega-3 fatty acids in human diets and higher fat equates to higher levels of omega-3s reaching the end consumer (Tocher, 2015). Lipid content is a highly polygenic trait, with many genes explaining a small fraction of the total genetic variation (Pena et al., 2016). In Atlantic salmon, lipid content has been reported to have a heritability of 0.18 to 0.2 (Kristjánsson et al., 2020; Tsai et al., 2015), so breeding for fat content in Atlantic salmon is possible.

Breeding for complex traits such as growth rate, disease resistance, and filet properties has been accelerated in recent years with the widespread adoption of genomic selection. This breeding strategy takes advantage of genome wide single nucleotide polymorphism (SNP) data to calculate genomic breeding values (GBV) for each individual based on the genotype and phenotype of their parents, enabling rational selection of breeding pairs (Meuwissen et al., 2001). Genomic selection in salmon has already been successfully applied; with breeding programs improving lice resistance and filet color faster and more reliably than traditional breeding programs (Ødegård et al., 2014). Genomic selection for fat content has been successfully applied to rainbow trout (Hu et al., 2020), however such methods have yet to be applied to fillet fat in Atlantic salmon.

Application of genomic selection to production traits traditionally does not consider the biology of the target trait, and rather weighs all SNPs equally to predict genomic effects (Meuwissen et al., 2001). However, biological knowledge about the trait of interest can be used to improve the power and accuracy of genomic predictions (de las Heras-Saldana et al., 2020; MacLeod et al., 2016). For this reason, it is useful to understand the underlying changes in molecular physiology occurring during genomic selection to prioritize SNPs for future rounds of genomic selection. In Atlantic salmon, lipid homeostasis is achieved by balancing dietary intake from the gut, *de-novo* synthesis in the body, and excretion through gut or biliary systems, which involves the coordinated action of thousands of genes. Liver plays a central role in lipid metabolism, controlling the flow of dietary lipids between different parts of the body through absorption and secretion of different lipoproteins to the circulatory system (Vance and Vance, 2008). Liver is therefore a logical place to analyze the effect of genomic selection for fat content on the molecular physiology of the fish.

In this study, we calculate GBVs for filet fat content in 184 fish based on genotype and phenotype data from a training set of 2487 fish. We sequenced the liver transcriptomes of all 184 fish and associate gene expression with GBV. Finally, we took the most significant GBV associated genes (padj < 10^-4^) together with a manually curated subset of known lipid related GBV associated genes and identify eQTLs for each (Figure 1a). Our aim is to improve our understanding of the genetic basis for differences in fat content by interrogating the link between GBVs and gene regulation in the liver.

**Figure 1:**
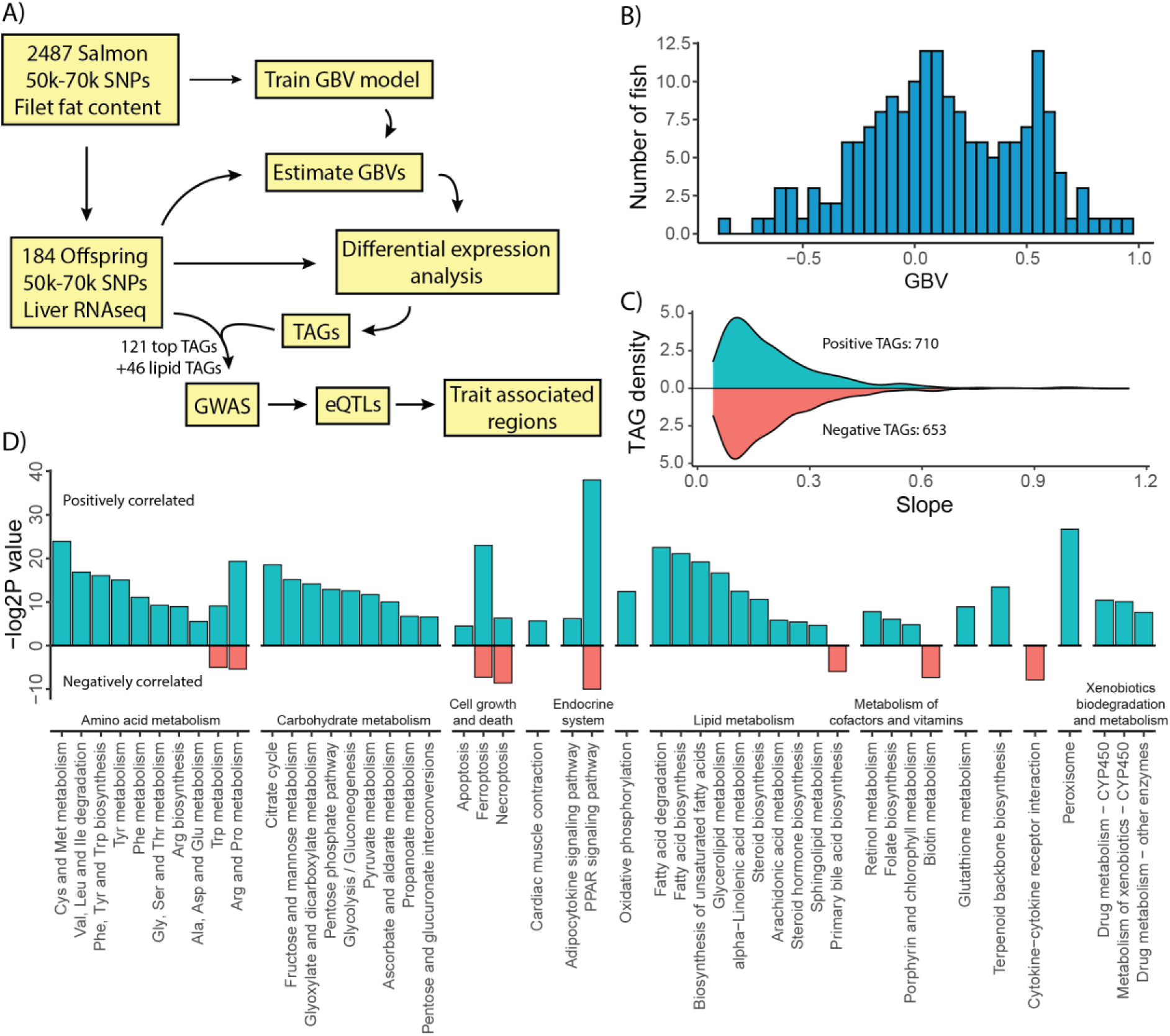
Correlation of GBV to gene expression. A) Flow chart of our analysis. B) Distribution of GBVs in the 184 salmon used for RNAseq analysis. C) Slope distribution among positively and negatively correlated TAGs. D) KEGG enrichment analysis of positively and negatively correlated TAGs. Only significantly enriched pathways (p < 0.05) are shown.

## 2 Methods

### 2.1 Fish and housing

A family experiment with Atlantic salmon was carried out at the fish laboratory, Norwegian University of Life Sciences (NMBU), Aas, Norway. The family experiment is explained in detail by Dvergedal *et al*. (Dvergedal et al., 2019). Broodstock from AquaGen’s breeding population (22 males and 23 females) were used to generate 23 families.

From the eyed egg stage until the start of the experiment, all families were communally reared in a single tank until the start of the experiment. When the fish reached 5-10 g, they were pit-tagged with a 2 x 12 mm unique glass tag (RFID Solutions, Hafrsfjord, Norway), and a fin-clip was collected for DNA-extraction and genotyping of a total of 2,300 fish. Fin clips (20 mg) were incubated in lysis buffer and treated with proteinase K (20 µg/ml) at 56 ℃ overnight. The following day, DNA was isolated from the lysate using the sbeadex livestock kit (LGC Genomics) according to the manufacturer’s protocol (Thermo Fisher Scientific) at Biobank AS (Hamar, Norway). The DNA concentration was measured using a Nanodrop 8000 (Thermo Fisher Scientific). All fish were genotyped using AquaGen’s custom Axiom®SNP (single-nucleotide polymorphism) genotyping array from Thermo Fisher Scientific (former Affymetrix) (San Diego, CA, USA). This SNP-chip contains 56,177 SNPs which were originally identified based on Illumina HiSeq reads (10-15x coverage) from 29 individuals from AquaGen’s breeding population. Genotyping was done at CIGENE (Aas, Norway). Genotypes were called from the raw data using the Axiom Power Tools software from Affymetrix. Individuals having a Dish-QC score below 0.82, and/or a call-rate below 0.97 were deleted from further analyses. *A priori* to the 12-day test, the parentage of each fish was established using genomic relationship likelihood for parentage assignment (Grashei et al., 2018), and families were allocated to tanks, 50 fish per tank, and 2 tanks per family. Except for nine tanks in which the number of fish varied between 42 and 54, due to some mortality before the start of the experiment or an increased number due to a counting mistake. The total number of fish was 2,281 and families were fed a fishmeal-based diet, as described in Dvergedal *et al*. (Dvergedal et al., 2019). The phenotypic data were registered individually for relative growth, as described by Dvergedal *et al*. (Dvergedal et al., 2019).

### 2.2 Estimating genomic breeding values (GBV) for fat content

Estimated genomic breeding values (GBV) for fillet fat content were predicted using a training data set consisting of 2487 genotyped (50-70k SNP chip) and phenotyped (fillet fat content) fish from two slaughter tests performed on the parental year-class 2014 (766 fish of the parental generation of the fish in the current study) and 2017 (1721 fish of the same generation as in the current study). Phenotypes for fat content were obtained using Norwegian quality cuts (NQC) from each fish that were subsequently frozen. The cut was thawed, skin and central bone removed, and homogenized for fat measurement on a NIR XDS machine. Fat content was predicted using a proprietary NQC model owned by Cargill. Average fat (standard deviation) was 12.96% (1.22%) and 17.41% (2.84%) for, respectively, year-classes 2014 and 2017. Additionally, 59 samples were analyzed by an independent lab (Eurofins) for validation. A linear genomic animal model (GBLUP) was used to obtain EBV:

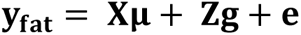

Where **y_fat_** is a vector of fat content phenotypes (standardized within each year-class), **μ** is a vector of the two year-class intercepts, 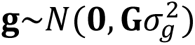 is a vector of polygenic effects (including both 2014 and 2017 year-class fish in the current study), **e** is a vector of random residuals, 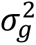 is the additive genetic variance, **G** = ρ**MM′** is the genomic relationship matrix, **M** is a centered genotype matrix (one row per individual and one column per locus), 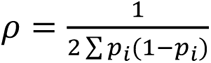 and *p_i_* is the allele frequency at locus *i*. The estimated genomic heritability (across the two year-classes) for fillet fat content was rather moderate (0.23 ± 0.03). SNP marker effects estimates were obtained as: 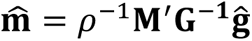 (Legarra et al., 2018). Using the estimated marker effects GBVs for the fish in the current study were predicted as:

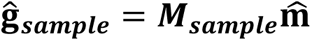

Where 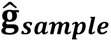 is a vector of GBVs (on the standardized scale) for fat content of fish in the current study and ***M_sample_*** is a (centered) genotype matrix of the same fish.

### 2.3 RNA extraction and transcriptomic sequencing

Four individual fish from each family were used for RNA isolation. RNA was extracted from liver of each individual fish using RNeasy Plus Universal Kit (Qiagen, Hilden, Germany), according to the manufacturer’s instructions. The concentration of RNA was determined by a Nanodrop 8000 (Thermo Fisher Scientific, Waltham, USA), and RNA integrity was examined by using a 2100 Bioanalyzer (Agilent Technologies, Santa Clara, USA). All RNA samples had RNA integrity (RIN) values higher than 8. Sequencing libraries were generated using TruSeq Stranded mRNA Library Prep Kit (Illumina, San Diego, USA) according to the manufacturer’s protocol. Libraries were delivered to Norwegian Sequencing Centre (Oslo, Norway) where all 184 samples were merged into one flow cell and sequenced using 100bp single-end mRNA sequencing (RNA-seq) on Illumina Hiseq 2500 (Illumina, San Diego, CA, USA).

Raw fastq file of reads sequences are publicly available on ArrayExpress under accession number E-MTAB-8305. Gene expression was quantified using the Salmon quasi-mapper version 0.13.1 (Patro et al., 2017) against the Atlantic salmon transcriptome (ICSASG_v2).

### 2.4 Trait association analysis

Trait associated genes (TAGs) were detected by correlating gene expression to GBV using edgeR (Robinson et al., 2009). Genes with low read counts, i.e. less than 0.5 count per million (CPM) in 50% of the samples, were removed and GBV was scaled (mean = 0 and standard deviation =1) prior to analysis. The linear regression model was

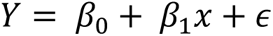

where Y is gene expression and x is scaled GBV value. Genes were classified as TAGs if the slope (β_1_) was significant, i.e. β_1_ ≠ 0 with false discover rate adjusted p value < 0.05. We also ran the analysis with family as a fixed effect factor in the model, however this only slightly changed the number of TAGs and did not influence the conclusion, so we use the simpler model in the analysis. KEGG enrichment analysis of the TAGs with positive and negative slope was performed with the kegga function in the limma R package (Ritchie et al., 2015). Translations between human readable gene names and NCBI RefSeq annotation gene ID’s can be found in File S1.

### 2.5 Genome-wide association analysis

To associate variation in TAGs with host genetics, a genome-wide association study was done using TAGs with an adjusted p-value <0.0001 (*n* = 121) and genes known to be associated with fat metabolism in the liver (*n* = 46) as response variables. The analysis was carried out by a linear mixed-model algorithm implemented in a genome-wide complex trait analysis (GCTA) (Yang et al., 2011). The leave one chromosome out option (--mlm-loco) was used, meaning that the chromosome harboring the SNP tested for was left out when building the genetic relationship matrix (GRM). The linear mixed model can be written as:

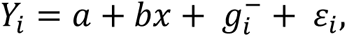

where *Y_i_* is one of the TAGs of individual *i*, *a* is the intercept, *b* is the fixed regression of the candidate SNP to be tested for association, *x* is the SNP genotype indicator variable coded as 0, 1, or 2, 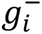 i is the random polygenic effect for individual 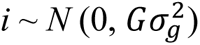 where **G** is the GRM and 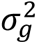 is the variance component for the polygenic effect, and *ε_i_* is the random residual. In this algorithm, 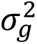 is re-estimated each time a chromosome is left out from the calculation of the GRM. The dataset was filtered, and individuals with < 10% missing genotypes were kept (*n* = 2279). Further, it was required that SNPs should have minor allele frequency (MAF) ≥ 1% and a call rate > 90%. After filtering, 54,200 SNPs could be included in the analysis. The level of significance for SNP was evaluated with a built-in likelihood-ratio test, and the threshold value for genome-wide significance was calculated using the Bonferroni correction (0.05/ 54200 = 9.23 × 10−7), corresponding to a -log10 *p*-value of 6.03.

### 2.6 Allele distribution of top SNPs for TAGs associated with lipid metabolism in liver

For TAGs with a significant genome-wide eQTL association to lipid metabolism in liver, the allele distribution for the top SNP (the SNP with the highest -log10 *p*-value) was examined with PLINK 2.00 (Chang et al., 2015) using the --recode option, which creates a new file after applying sample/variant filters and the --extract option to create a file with the top SNP of interest.

### 2.7 Co-expression network analysis

We assembled an independent RNA-seq expression data set comprising 112 liver samples spanning different diets and life stages in fresh water (Gillard et al., 2018). Raw RNA-Seq data can be found in the European Nucleotide Archive (ENA) under project accession no. PRJEB24480. Read counts for Atlantic salmon genes (NCBI: GCF_000233375.1_ICSASG_v2) were estimated using Salmon (Patro et al., 2017). Code and data available at https://gitlab.com/garethgillard/megaLiverRNA. 27,786 genes with at least 10 mapped reads in more than 10 samples were retained for further analysis. Read counts were normalized using the varianceStabilizingTransformation-function from the R package DESeq2 (Love et al., 2014).

For co-expression network inference, we used the Weighted Gene Co-expression Network Analysis (WGCNA) R package (Langfelder and Horvath, 2008) and the function blockwiseModules with the bicor correlation measure and parameters *power = 3 (scale free topology fit with an R2 of 0.8), maxBlockSize = 30000, networkType = “signed”, TOMType = “signed”, corType = “bicor”, maxPOutliers = 0.05, replaceMissingAdjacencies = TRUE, pamStage = F, deepSplit = 1, minModuleSize = 5, minKMEtoStay = 0.2, minCoreKME = 0.2, minCoreKMESize = 2, reassignThreshold = 0 and mergeCutHeight = 0.2*. A hard correlation threshold of 0.5 was used to visualize the network in Cytoscape (https://cytoscape.org).

## 3 Results

### 3.1 Correlations between liver gene expression levels and GBV

We first calculated GBV for the 184 fish used in this study based on a training set of 2487 fish (Figure 1a and b). RNA sequencing and trait association analysis revealed a total of 710 TAGs (padj < 0.05) positively correlated to GBV and 653 TAGs negatively correlated to GBV (Figure 1c, File S2). By comparing the number of TAGs to total genes within each KEGG pathway, we found that the positive TAGs were enriched (p < 0.05) in 45 KEGG pathways (Figure 1d, File S3). Many of the most significantly enriched KEGG pathways were related to lipid metabolism including “*PPAR signaling*”, “*fatty acid degradation”,* “*fatty acid biosynthesis*”, “*biosynthesis of unsaturated fatty acids*” and “*glycerolipid metabolism*” (Figure 1d). In addition, pathways related to amino acid metabolism and energy metabolism were enriched for positively correlated TAGs (Figure 1d). Only eight pathways were found enriched (p < 0.05) for the negatively correlated TAGs. This included the lipid metabolism pathway *“primary bile acid synthesis”* which produces bile acids from cholesterol that aid in lipid solubilization in the intestine.

### 3.2 Regulation of lipid metabolism genes

To further dissect the link between lipid metabolism and GBV for fat content, we performed an in-depth analyses of a manually annotated set of genes involved in lipid metabolism in salmon (Gillard et al., 2018). We found that the TAG list was clearly enriched for lipid genes (fisher’s p-value 2.2e-16, odds ratio 4.92) with 45 (6.3%) lipid genes found in the 710 TAGs positively correlated to GBV, and 10 lipid genes (1.5%) found in the 653 TAGs negatively correlated with GBV (Figure 2a). Intriguingly, the GBV-associated lipid genes covered genes involved in many enzymatic steps in the cholesterol biosynthesis pathway including the enzyme controlling the rate limiting step of cholesterol biosynthesis, *hmgcrab*. Other lipid metabolism enzyme-encoding genes associated to fat content GBV included fatty acid synthase (*fasn-b),* all three polyunsaturated fatty acid desaturases *(fads2d5 fads2d6a, and fads2d6b)* (figure 2e), and several lipid and fatty acid transporters (*fatp2f*, *fabp7b*, *ldlrab-a*, and *apobc)* (Figure 2c).

**Figure 2:**
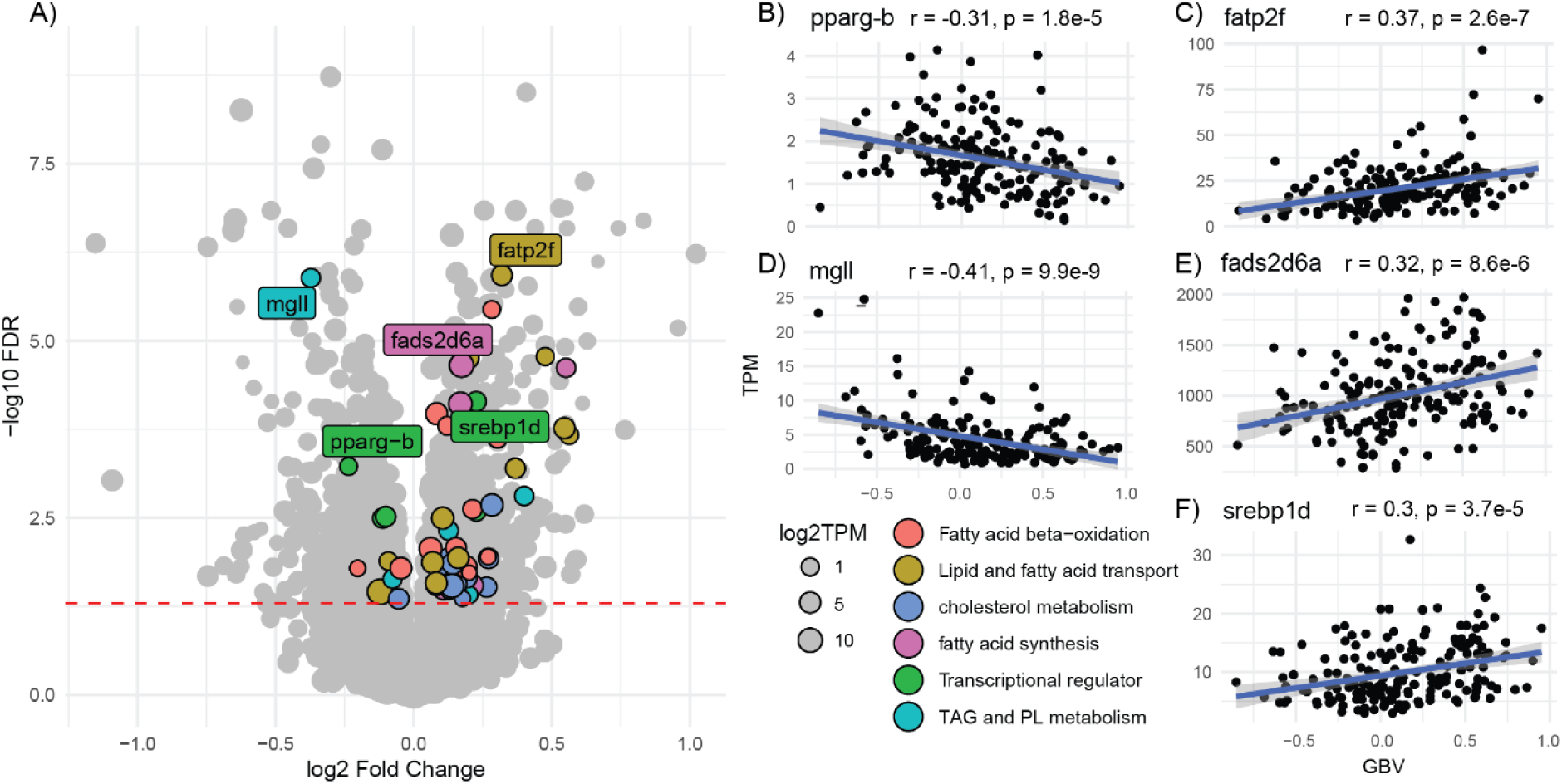
Correlation between GBV and lipid metabolism gene expression. A) Volcano plot of the regression results between gene expression and GBV. Genes involved in lipid metabolism are coloured and important lipid metabolism genes are labelled. Size corresponds to mean log2TPM values. The dashed red line indicates the padj < 0.05 cutoff. Correlation of GBV and gene expression are shown on the right for *pparg-b* (B), *fatp2f* (C), and *mgll* (D), *fads2d6a* (E), and *srebp1d* (F).

We also found several lipid regulatory genes positively correlated with GBV (Figure 2), including all three isoforms of SREPB1 (*srebp1b, srebp1c, srebp1d)* (Figure 2f), known regulators of lipid biosynthesis (Shimano and Sato, 2017). Lipid metabolism genes negatively correlated to GBV included the important lipid oxidation regulator *pparg-b* (figure 2b) as well as both copies of farnesoid x receptor (*fxr-a* and *fxr-b*), which plays a key role in hepatic triglyceride homeostasis and is involved in suppression of lipogenesis (Jiao et al., 2015). The lipid gene that was most negatively associated with GBV was monoacylglycerol lipase (*mgll*) which is involved glycerolipid breakdown.

### 3.3 eQTL analyses highlights several trans-acting loci impacting many genes

To explore the genetic architecture of gene expression differences between fish with high and low GBV, we used linear regressions to identify genetic variants associated gene expression levels (File S4). In total 167 genes associated with GBV fat were examined, which included the top 121 significant TAGs (padj < 0.0001), as well as 46 significant TAGs from lipid pathways (padj < 0.05) (Figure 2a). Genes that were not annotated on a chromosome (i.e belonged to a smaller unplaced scaffold) were discarded from this analysis.

In total, 71 genes had at least one significant eQTL (genome wide significance level, p < 10^-6^), with a mean number of 1.4 eQTLs per gene (Figure 3a). Dissecting the positions of eQTL signals relative to these genes showed that 21 genes had a significant association in cis (no more than 10 Mbp from the gene). Considering only “top associations” for each gene reveals a clear tendency for eQTL-associations to variants on other chromosomes (i.e. trans associations) (Figure 3b). Seven of the genes with significant eQTL’s were known lipid metabolism genes; two of which are monoglyceride lipase (*mgll*) and fatty acid desaturase 2-like (*fads2d6b*) (Figure 3c-f).

**Figure 3:**
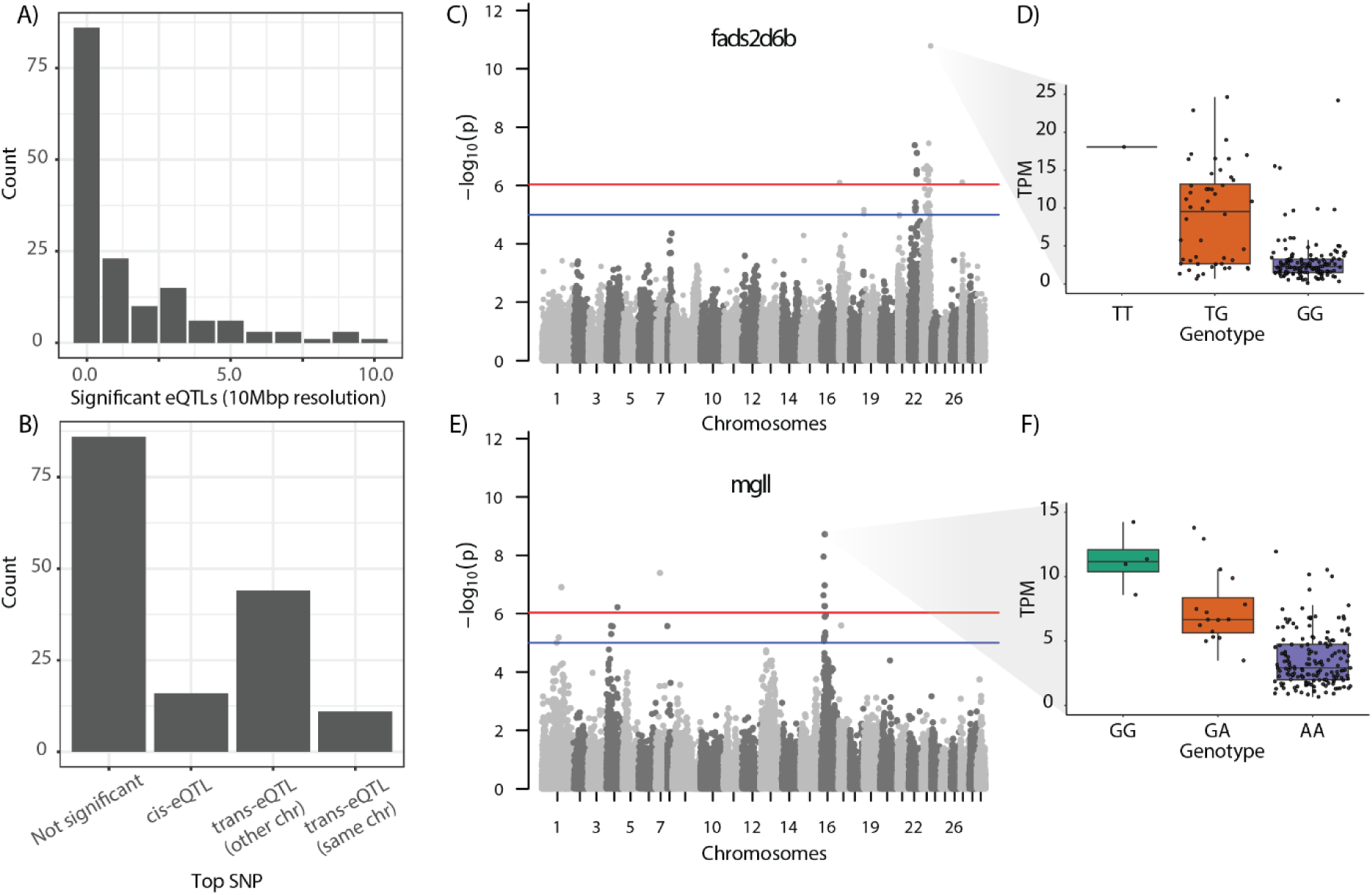
GWAS of top GBV TAGs and selected lipid metabolism genes. A) Number of significant (p < 10^-6^) eQTLs per gene. B) Number of genes with the top eQTL in cis (< 10 MB from gene), trans (different chromosome), or trans (same chromosome). C) Manhattan plot of SNPs associated with *fads2d6b* gene expression. Significance cutoffs are indicated by red (strict) and blue (relaxed) lines. D) Genotype distribution of the top SNP for *fads2d6b*. E and F) Same as C and D for the gene *mgll*.

Since the genes tested for eQTLs were associated with differences in one quite specific molecular trait (i.e. lipid metabolism), it is plausible that a few top regulators of key lipid-metabolism pathways could impact the expression of many of the genes in our study. The high numbers of trans eQTL signals (Figure 3b) supports this idea. Hence, to test if any chromosomal regions showed sign of harboring such major regulators, we analyzed the distribution of top trans-eQTL associations across chromosomes. Using a relaxed p-value cutoff for significant associations (p < 10^-4^), two chromosomes (3 and 6) were clearly enriched for top trans-eQTLs relative to the total number of SNPs on each chromosome (Figure 4a). It is worth noting, that the enrichment signal on chromosome 6 dropped rapidly as we increased p-value cutoff stringency and was diminished at p < 10^-6^.

**Figure 4:**
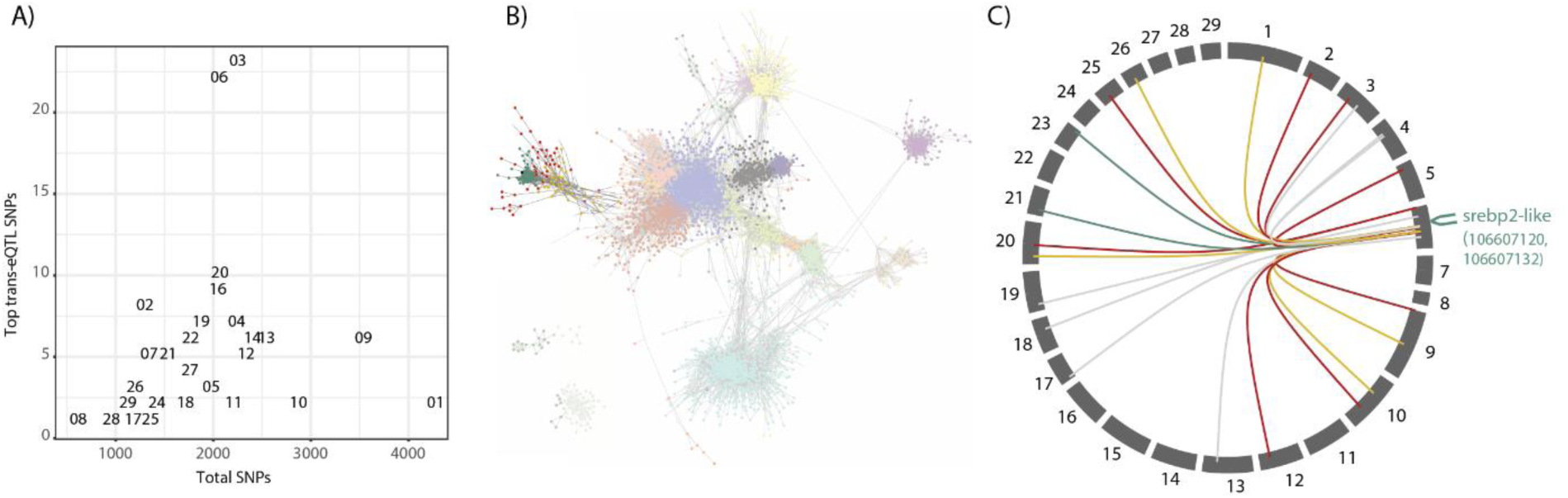
Trans eQTL and gene co-expression. A) Relationship between total number of SNPs for each chromosome and the number of genes having top trans-eQTL signals on each chromosome. B) Co-expression network from liver RNAseq data highlighting three co-expression modules which contain genes with top trans eQTL signals (p < 10^-4^) on chromosome 6. C) Circos plot showing the top trans-eQTL links between trait associated genes and chromosome 6. Colors reflect the co-expression module that the trait associated genes belong to. We have indicated the position of a major lipid metabolism transcription factor (srebp2) on chromosome 6.

Next, we performed an in-depth analysis of the eQTL signals on chromosome 3 and 6, with specific focus on lipid-metabolism genes. We hypothesized that if regulators of lipid-metabolism pathways were located on these chromosomes we would find clusters of trans-eQTLs for genes that are co-regulated in liver. We therefore first used a large gene expression dataset of 112 liver samples to estimate co-expression modules (i.e. genes whereby their expression correlates across different samples) (Figure 4b). Genes within these co-expression modules are predicted to share transcriptional regulators. We then associated genes with trans-eQTLs on chromsomes 6 and 3 to specific co-expression modules. Even though the trans-eQTL enrichment on chromosome 3 contained many more highly significant associations compared to chromosome 6 (Figure S1), there was no obvious clustering of trans-eQTL signals among co-expressed genes on this chromosome. However, the central region (35-50 Mbp) of chromosome 6 displayed a clear clustering of trans-eQTL signals to co-expressed genes. Although these genes belonged to three different co-expression modules, these three modules were virtually identical in the co-expression network (Figure 4b). Interestingly, this genomic region harbors two copies of srebp-2-like genes, known to regulate various aspects of lipid-metabolism in vertebrates, which also belong to one of these three co-expression modules.

## 4 Discussion

Our results clearly demonstrate that genomic selection for high muscle fat content in salmon drives changes in expression of genes involved in lipid metabolism in the liver. Variation in muscle lipid content among fish could be due to differences in 1) uptake of lipids from the diet, 2) *de-novo* lipogenesis in the body, 3) lipid transport and deposition between different part of the body, 4) lipid degradation by beta-oxidation and efflux through bile synthesis, or 5) a combination. Our results have shown that GBV was positively associated with gene expression in lipid transport (*fatp2f*, *fabp7b*), *de novo* lipogenesis (*fas*), fatty acid desaturation (*fads2d5*, *fads2d6a*, *fads2d6b*, and *fads2d6c*), cholesterol biosynthesis (*hmgcrab*), and negatively associated with genes in glycerolipid breakdown (*mgll*), beta-oxidation (*pparg-b*) and bile acid synthesis (*fxr*) in liver. This suggests that higher deposition of lipids in high GBV salmon was most likely due to a combination of reduced glycerolipid breakdown, elevated lipid synthesis, and elevated lipid transport. Both endogenous and exogeneous lipids in liver are packaged into very-low-density lipoproteins (VLDL) which are secreted into the vasculature. Lipids in VLDL are taken up by the peripheral tissues such as muscle, and the leftover lipoproteins (low-density lipoproteins, LDL) are taken back by liver through the LDL receptor (LDLR) (Wang, 2007). Most of the lipoproteins including VLDL and LDL contain a copy of apolipoprotein B (APOB), an essential component for its structure (Elovson et al., 1988). Our study has found GBV to be positively correlated to *ldlrab-a*, and *apobc*. This could suggest high GBV was associated with increased amount and turnover of lipoproteins in the bloodstream. Additionally, we have also identified two *fxr* genes which were negatively correelated to GBV. Since fxr is a key regulator and positively correlated to bile salt synthesis in fish (Wen et al., 2021), this suggests that high GBV fish has decreased lipid excretion through bile salt production pathway. Under this scenario, a higher proportion of newly synthesized saturated and monounsaturated fatty acid, and diet derived polyunsaturated fatty acid containing triacylglycerol are exported as lipoproteins from the liver and deposited in peripheral tissues including muscle where the fat phenotype was measured.

We find expression of key genes in lipid biosynthesis pathways, including *fas*, *fads2d5*, *fads2d6a*, *fads2d6b*, and *fads2d6c* were positively correlated to GBV. This is likely due to higher expression of sterol regulatory element-binding protein 1 (SREBP1), which is the key positive regulator of fatty acid *de-novo* synthesis and LC-PUFA synthesis in salmon (Carmona-Antoñanzas et al., 2014; Minghetti et al., 2011). Other genes of LC-PUFA synthesis, the *elovl2* and *elovl5* genes, were not correlated to GBV. This could be because these genes are not controlled by SREBP1 transcription factor in salmon (Datsomor et al., 2019). A similar study using high and low muscle fat lines of rainbow trout also found higher expression of lipogenic genes in high fat lines and hypothesized that this was due to a more active target of rapamycin (TOR) signalling pathway (Skiba-Cassy et al., 2009). Fat and lean rainbow trout lines displayed a similar ratio of phosphorylated and native TOR, however fat lines had significantly higher levels of TOR protein. We did not find TOR mRNA to be associated with GBV of muscle lipid levels in Atlantic salmon, however TOR protein abundance could be regulated at the posttranslational level. In addition, we found that ATP-citrate lyase (*acyl*, geneID:106589258) gene expression was positively associated to GBV, which agrees with the study in rainbow trout (Skiba-Cassy et al., 2009). ACYL acts as a metabolic switch linking glucose and lipid metabolism that diverts citrate from the TCA cycle into lipogenesis by converting it to acetyl-CoA and oxaloacetate in the cytosol. This enables elevated lipogenesis by increasing the available pool of acetyl-CoA to be converted to fatty acids by FAS.

Our eQTL analysis revealed an unexpectedly high number of trans-eQTLs on chromosome 6 associated to co-regulated genes that were TAGs in our analysis. Moreover, these chromosome trans-eQTL associations were mostly originating from a smaller region around 50 Mbp, pointing to a potential common transcription factor. Two of the genes in this region are paralogs of SREBP2-like, a known regulator of lipid metabolism. Srebp2 is a key transcriptional regulator controlling cholesterol metabolism in fish and mammals (Carmona-Antoñanzas et al., 2014; Madison, 2016), and our study has found many positively correlated TAGs involved in *de-novo* cholesterol synthesis including *hmgcrab*. Additionally, positively correlated *acyl* suggests an increased acetyl-CoA pool. Although we did not find *srebp2* to be associated with GBV in our analysis, SREBP is known to be highly regulated post-transcriptionally through interactions with SREBP cleavage-activating protein (SCAP). SCAP forms a complex with SREBP and facilitates the cleavage of SREBPs by site-1 protease, thereby releasing active

NH2-terminal fragments from the ER membrane to nucleus, activating gene expression (Nohturfft et al., 1998). Therefore, this lack of association could be explained by variation in *srebp2* protein structure resulting in increasing SCAP or Site-1 protease binding activity without influencing *srebp2* expression itself. Alternatively, it may be that the trans-eQTL signal cluster on chromosome 6 is driven by another, so far unknown regulator of lipid metabolism on chromosome 6.

Although lipid synthesis and transport in the liver contributes to lipid content in the muscle, our results only tell part of the story. Since there is considerable variation in fat deposition and turnover in salmon (Dvergedal et al., 2020; Dvergedal et al., 2019) variation in the regulation of lipid metabolism in the muscle must also be a large contributor to the high fat phenotype. High muscle fat has previously been associated with downregulation of genes related to lipid catabolism and upregulation of genes associated to glycogenolysis (Horn et al., 2019), which may signal a transition in how fish utilize energy stores. Additionally, differences in hormonal signalling between the brain, adipose, and muscle tissues could contribute to the high fat phenotype. To further improve our understanding of what makes a salmon fat, future studies need to address these aspects of the salmon molecular physiology.

## 5 Conclusion

We demonstrate that genomic selection using estimated breeding values for fat content drives changes in lipid metabolism in Atlantic salmon. Fish with high GBV for muscle fat content had overall higher gene expression in lipid biosynthesis and transport pathways and lower expression of genes involved in glycerolipid breakdown. This is important validation for genomic selection as a strategy to improve lipid content and the results could be used to prioritize SNPs in future estimates of breeding values.

## Ethics approval and consent to participate

The experiment used phenotypic data, which were collected from a family experiment with Atlantic salmon carried out at the fish laboratory, Norwegian University of Life Sciences (NMBU), Aas, Norway, following the laws and regulations for experiments on live animals in EU (Directive 2010/637EU) and Norway (FOR-2015-06-18-761). The experiment was approved by the Norwegian Food Safety Authority (FOTS ID 11676).

## Data availability

The genotypic data are owned by AquaGen AS, used under license for this study, and not publicly available. Phenotypic data can be made available on request. All RNAseq data is publicly available on ArrayExpress under accession number E-MTAB-8305.

## Acknowledgements

We thank Bjørn Reidar Hansen, Harald Støkken and Bjørn Frode Eriksen for help and assistance at the fish laboratory, and Ricardo Tavares Benicio, Ragnhild Ånestad, Milena Bjelanovic, Jon Øvrum Hansen, Mathabela Nelson, Shomorin Oluwaseun George and Ingrid Marie Håkenåsen for their help during feed production, experiment, and sampling. A thank goes to all that contributed during sampling of experimental material.

## Supplementary data

**Figure S1: Trans eQTL links on chromosome 3.** Circos plot showing the top trans-eQTL links between trait associated genes and chromosome 3. Colors reflect the co-expression module that the trait associated genes belong to from Figure 4b.

**File S1: Human readable gene name to NCBI id translations.** List of human readable gene names of lipid genes used in our analysis and their associated NCBI refseq identifiers. Column 1 - NCBI gene ID, column 2 – human readable gene name, column 3 – gene description.

**File S2: Trait associated genes for fat content in Atlantic salmon.** List of all salmon genes significantly associated to GBV. Column 1 – Gene ID, column 2 – log2 fold change, column 3 – adjusted p-value, column 4 – gene name, column 5 – gene product.

**File S3: Enriched KEGG pathways for GBV associated genes.** List of all significantly (p < 0.05) enriched KEGG pathways among GBV associated genes. Column 1 – pathway name, column 2 – pathway ID, column 3 – type of GBV correlation for genes in pathway, column 4 – number genes in pathway, column 5 – number of TAGs in pathway, column 6 – enrichment p-value.

**File S4: eQTL results of selected TAGs.** List of SNPs significantly associated to TAG expression (p < 10^-6^). Column 1 – NCBI gene ID of TAG, column 2 – Chromosome containing SNP, column 3 – SNP identifier, column 4 – Physical position of SNP, column 5 – Reference allele, column 6 – Second allele, column 7 – Frequency of the reference allele, column 8 – SNP effect, column 9 – Standard error, column 10 – p-value.

## References

Carmona-Antoñanzas, G., Tocher, D.R., Martinez-Rubio, L., Leaver, M.J., 2014. Conservation of lipid metabolic gene transcriptional regulatory networks in fish and mammals. Gene 534(1), 1–9. 10.1016/j.gene.2013.10.040

Chang, C.C., Chow, C.C., Tellier, L.C., Vattikuti, S., Purcell, S.M., Lee, J.J., 2015. Second-generation PLINK: rising to the challenge of larger and richer datasets. GigaScience 4(1). 10.1186/s13742-015-0047-8

Datsomor, A.K., Zic, N., Li, K., Olsen, R.E., Jin, Y., Vik, J.O., Edvardsen, R.B., Grammes, F., Wargelius, A., Winge, P., 2019. CRISPR/Cas9-mediated ablation of elovl2 in Atlantic salmon (Salmo salar L.) inhibits elongation of polyunsaturated fatty acids and induces Srebp-1 and target genes. Scientific Reports 9(1). 10.1038/s41598-019-43862-8

de las Heras-Saldana, S., Lopez, B.I., Moghaddar, N., Park, W., Park, J.-e., Chung, K.Y., Lim, D., Lee, S.H., Shin, D., van der Werf, J.H.J., 2020. Use of gene expression and whole-genome sequence information to improve the accuracy of genomic prediction for carcass traits in Hanwoo cattle. Genet. Sel. Evol. 52(1), 54. 10.1186/s12711-020-00574-2

Dvergedal, H., Harvey, T.N., Jin, Y., Ødegård, J., Grønvold, L., Sandve, S.R., Våge, D.I., Moen, T., Klemetsdal, G., 2020. Genomic regions and signaling pathways associated with indicator traits for feed efficiency in juvenile Atlantic salmon (Salmo salar). Genet. Sel. Evol. 52(1), 66. 10.1186/s12711-020-00587-x

Dvergedal, H., Ødegård, J., Øverland, M., Mydland, L.T., Klemetsdal, G., 2019. Selection for feed efficiency in Atlantic salmon using individual indicator traits based on stable isotope profiling. Genet. Sel. Evol. 51(1), 13. 10.1186/s12711-019-0455-9

Elovson, J., Chatterton, J.E., Bell, G.T., Schumaker, V.N., Reuben, M.A., Puppione, D.L., Reeve, J.R., Young, N.L., 1988. Plasma very low density lipoproteins contain a single molecule of apolipoprotein B. J. Lipid Res. 29(11), 1461–1473. 10.1016/S0022-2275(20)38425-X

Gillard, G., Harvey, T.N., Gjuvsland, A., Jin, Y., Thomassen, M., Lien, S., Leaver, M., Torgersen, J.S., Hvidsten, T.R., Vik, J.O., Sandve, S.R., 2018. Life-stage-associated remodelling of lipid metabolism regulation in Atlantic salmon. Mol. Ecol. 27(5), 1200–1213. 10.1111/mec.14533

Grashei, K.E., Ødegård, J., Meuwissen, T.H.E., 2018. Using genomic relationship likelihood for parentage assignment. Genet. Sel. Evol. 50(1), 26. 10.1186/s12711-018-0397-7

Horn, S.S., Sonesson, A.K., Krasnov, A., Moghadam, H., Hillestad, B., Meuwissen, T.H.E., Ruyter, B., 2019. Individual differences in EPA and DHA content of Atlantic salmon are associated with gene expression of key metabolic processes. Scientific Reports 9(1), 3889. 10.1038/s41598-019-40391-2

Hu, G., Gu, W., Cheng, L., Wang, B., 2020. Genomic prediction and variance component estimation for carcass fat content in rainbow trout using SNP markers. J. World Aquacult. Soc. 51(2), 501–511. 10.1111/jwas.12677

Jiao, Y., Lu, Y., Li, X.-y., 2015. Farnesoid X receptor: a master regulator of hepatic triglyceride and glucose homeostasis. Acta Pharmacologica Sinica 36(1), 44–50. 10.1038/aps.2014.116

Kristjánsson, Ó.H., Gjerde, B., Ødegård, J., Lillehammer, M., 2020. Quantitative Genetics of Growth Rate and Filet Quality Traits in Atlantic Salmon Inferred From a Longitudinal Bayesian Model for the Left-Censored Gaussian Trait Growth Rate. Frontiers in Genetics 11. 10.3389/fgene.2020.573265

Langfelder, P., Horvath, S., 2008. WGCNA: an R package for weighted correlation network analysis. BMC Bioinformatics 9(1), 559. 10.1186/1471-2105-9-559

Legarra, A., Ricard, A., Varona, L., 2018. GWAS by GBLUP: Single and Multimarker EMMAX and Bayes Factors, with an Example in Detection of a Major Gene for Horse Gait. G3 Genes|Genomes|Genetics 8(7), 2301–2308. 10.1534/g3.118.200336

Love, M.I., Huber, W., Anders, S., 2014. Moderated estimation of fold change and dispersion for RNA-seq data with DESeq2. Genome Biology 15(12), 550. 10.1186/s13059-014-0550-8

MacLeod, I.M., Bowman, P.J., Vander Jagt, C.J., Haile-Mariam, M., Kemper, K.E., Chamberlain, A.J., Schrooten, C., Hayes, B.J., Goddard, M.E., 2016. Exploiting biological priors and sequence variants enhances QTL discovery and genomic prediction of complex traits. BMC Genomics 17(1), 144. 10.1186/s12864-016-2443-6

Madison, B.B., 2016. Srebp2: A master regulator of sterol and fatty acid synthesis1. J. Lipid Res. 57(3), 333–335. 10.1194/jlr.C066712

Meuwissen, T.H.E., Hayes, B.J., Goddard, M.E., 2001. Prediction of Total Genetic Value Using Genome-Wide Dense Marker Maps. Genetics 157(4), 1819–1829. 10.1093/genetics/157.4.1819

Minghetti, M., Leaver, M.J., Tocher, D.R., 2011. Transcriptional control mechanisms of genes of lipid and fatty acid metabolism in the Atlantic salmon (Salmo salar L.) established cell line, SHK-1. Biochim. Biophys. Acta 1811(3), 194–202. 10.1016/j.bbalip.2010.12.008

Nohturfft, A., Brown, M.S., Goldstein, J.L., 1998. Sterols regulate processing of carbohydrate chains of wild-type SREBP cleavage-activating protein (SCAP), but not sterol-resistant mutants Y298C or D443N. Proceedings of the National Academy of Sciences 95(22), 12848–12853. 10.1073/pnas.95.22.12848

Patro, R., Duggal, G., Love, M.I., Irizarry, R.A., Kingsford, C., 2017. Salmon provides fast and bias-aware quantification of transcript expression. Nat. Methods 14(4), 417–419. 10.1038/nmeth.4197

Pena, R.N., Ros-Freixedes, R., Tor, M., Estany, J., 2016. Genetic Marker Discovery in Complex Traits: A Field Example on Fat Content and Composition in Pigs. Int J Mol Sci 17(12). 10.3390/ijms17122100

Ritchie, M.E., Phipson, B., Wu, D., Hu, Y., Law, C.W., Shi, W., Smyth, G.K., 2015. Limma powers differential expression analyses for RNA-sequencing and microarray studies. Nucleic Acids Res. 43(7), e47–e47. 10.1093/nar/gkv007

Robinson, M.D., McCarthy, D.J., Smyth, G.K., 2009. edgeR: a Bioconductor package for differential expression analysis of digital gene expression data. Bioinformatics 26(1), 139–140. 10.1093/bioinformatics/btp616

Shimano, H., Sato, R., 2017. SREBP-regulated lipid metabolism: convergent physiology — divergent pathophysiology. Nature Reviews Endocrinology 13(12), 710–730. 10.1038/nrendo.2017.91

Skiba-Cassy, S., Lansard, M., Panserat, S., Médale, F., 2009. Rainbow trout genetically selected for greater muscle fat content display increased activation of liver TOR signaling and lipogenic gene expression. American Journal of Physiology-Regulatory, Integrative and Comparative Physiology 297(5), R1421–R1429. 10.1152/ajpregu.00312.2009

Tocher, D.R., 2015. Omega-3 long-chain polyunsaturated fatty acids and aquaculture in perspective. Aquaculture 449, 94–107. 10.1016/j.aquaculture.2015.01.010

Tsai, H.Y., Hamilton, A., Guy, D.R., Tinch, A.E., Bishop, S.C., Houston, R.D., 2015. The genetic architecture of growth and fillet traits in farmed Atlantic salmon (Salmo salar). BMC Genet. 16, 51. 10.1186/s12863-015-0215-y

Vance, D.E., Vance, J.E., 2008. CHAPTER 8 - Phospholipid biosynthesis in eukaryotes, in: Vance, D.E., Vance, J.E. (Eds.), Biochemistry of Lipids, Lipoproteins and Membranes (Fifth Edition). Elsevier, San Diego, pp. 213–244.

Wang, D.Q.H., 2007. Regulation of Intestinal Cholesterol Absorption. Annu. Rev. Physiol. 69(1), 221–248. 10.1146/annurev.physiol.69.031905.160725

Wen, J., Mercado, G.P., Volland, A., Doden, H.L., Lickwar, C.R., Crooks, T., Kakiyama, G., Kelly, C., Cocchiaro, J.L., Ridlon, J.M., Rawls, J.F., 2021. Fxr signaling and microbial metabolism of bile salts in the zebrafish intestine. Science Advances 7(30), eabg1371. 10.1126/sciadv.abg1371

Yang, J., Lee, S.H., Goddard, M.E., Visscher, P.M., 2011. GCTA: A Tool for Genome-wide Complex Trait Analysis. The American Journal of Human Genetics 88(1), 76–82. 10.1016/j.ajhg.2010.11.011

Ødegård, J., Moen, T., Santi, N., Korsvoll, S.A., Kjøglum, S., Meuwissen, T.H.E., 2014. Genomic prediction in an admixed population of Atlantic salmon (Salmo salar). Frontiers in Genetics 5. 10.3389/fgene.2014.00402

