## Supplementary figures and images for "Linking genomic prediction for muscle fat content in Atlantic salmon to underlying changes in lipid metabolism regulation"

### Figure S1

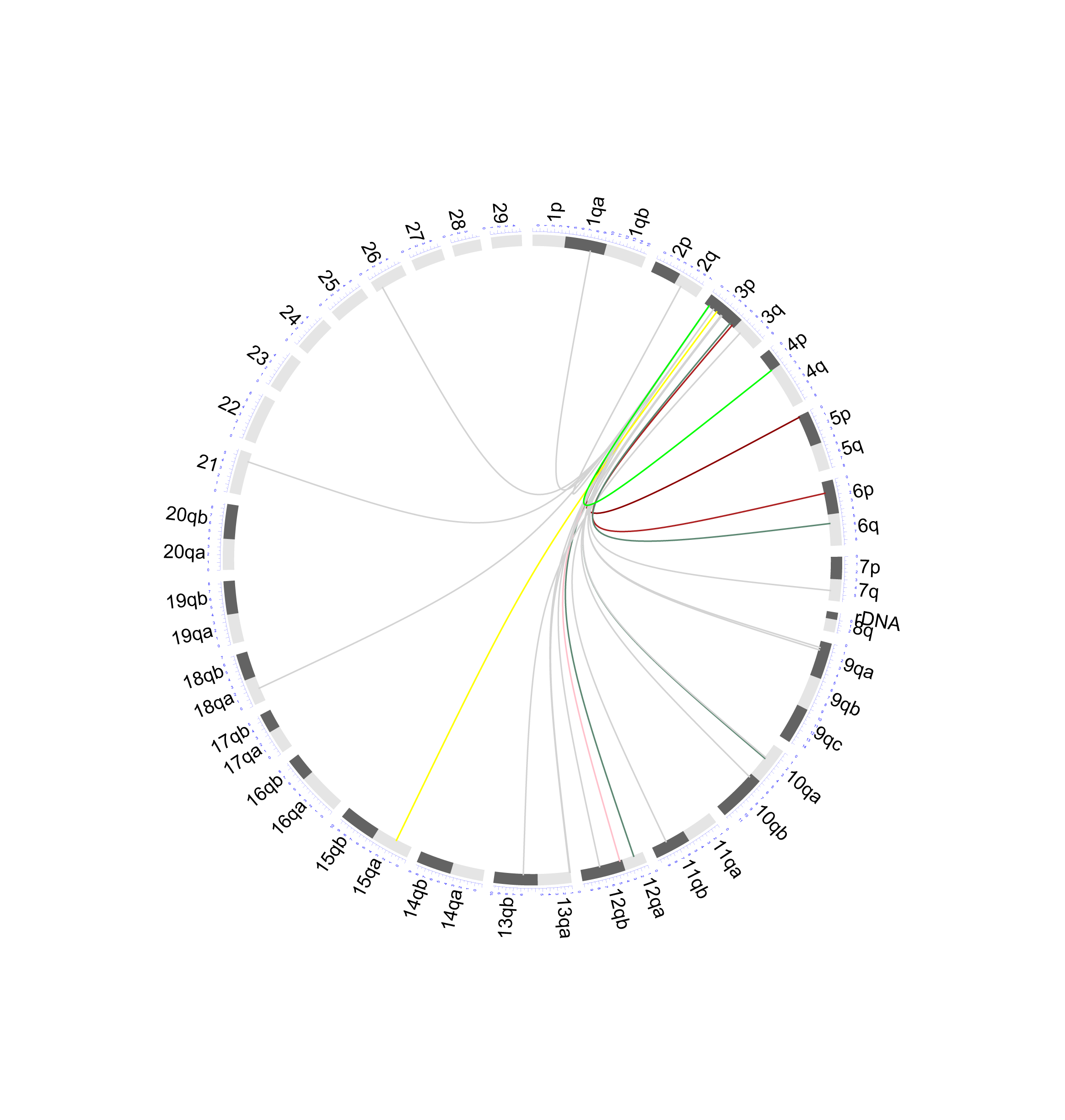
